# Modulation of cortico-muscular coupling associated with split-belt locomotor adaptation

**DOI:** 10.1101/2024.09.09.612154

**Authors:** Atsushi Oshima, Hikaru Yokoyama, Naotsugu Kaneko, Ryogo Takahashi, Ken Takiyama, Kimitaka Nakazawa

## Abstract

Humans can adjust their walking patterns according to the demands of their internal and external environments, referred to as locomotor adaptation. Although significant functional coupling (i.e. cortico-muscular coherence [CMC]) has been shown between cortical and lower-limb muscle activity during steady-state walking, little is known about CMC in locomotor adaptation. Therefore, we investigated the adaptation-dependent modulation of the CMC between the sensorimotor region and the tibialis anterior muscle using a split-belt locomotor adaptation paradigm that can impose an asymmetric perturbation. We hypothesized that the CMC would temporarily decrease after exposure to the asymmetric perturbation and removal of the perturbation because of a mismatch between the predicted and actual sensory feedback. We also hypothesized that the CMC would increase as adaptation and de-adaptation to perturbation progressed because the motor system could become able to predict sensory feedback. Our findings revealed that the CMC temporarily decreased after exposure to and removal of the perturbation. Moreover, the CMC increased with adaptation and de-adaptation to perturbation. Although these results depend on the leg, frequency bands, and gait phases, they partially support our hypothesis. These findings suggest that flexible updating of cortico-muscular coupling in the motor system is a key mechanism underlying locomotor adaptation in humans. The results from our study on healthy young individuals contribute to the understanding of neuromuscular control of gait and provide valuable insight for optimising gait rehabilitation.

**Key points:**

- Locomotor adaptation plays a crucial role in our daily activities and gait rehabilitation.
- Although knowledge regarding the brain and muscle activities associated with locomotor adaptation has been accumulated, little is known about the functional coupling of the brain and muscle activities.
- Using high-density EEG and lower limb EMG, we demonstrated the modulation of cortico-muscular coherence between the sensorimotor region and the tibialis anterior muscle with adaptation and de-adaptation during a split-belt treadmill walking paradigm.
- Our findings suggest that flexible updating of cortico-muscular coupling in the motor system underlies locomotor adaptation in humans.
- Understanding the human brain’s control of muscles during split-belt locomotor adaptation will deepen our knowledge of neuromuscular control of gait and provide valuable insights for gait rehabilitation.

## Introduction

Humans can adjust their walking patterns according to the demands of their internal and external environments (e.g. body condition and walking surfaces), referred to as locomotor adaptation. Locomotor adaptation is thought to be driven by a mismatch between the predicted and actual sensory feedback (Kitago & Krakauer, 2013; Rothwell *et al*., 2021). Adaptability plays a crucial role in daily activities and gait rehabilitation (Bastian, 2008). Thus, investigating the neural mechanisms underlying locomotor adaptation is important to advance our understanding of the neural control of human walking and gait recovery in patients with neurological disorders.

A commonly used method for investigating locomotor adaptation is a split-belt treadmill walking paradigm (Reisman *et al*., 2005; Choi & Bastian, 2007; Mawase *et al*., 2013; Severini & Zych, 2022; Jacobsen & Ferris, 2023). This treadmill has two belts controlled independently, providing a novel perturbation in which walking speeds differ between the right and left legs (i.e. asymmetry perturbation). When this perturbation is imposed, walking patterns become asymmetrical; however, they gradually return to symmetry within several minutes (i.e. adaptation) (Reisman *et al*., 2005). Adaptation is also characterised by the re-presence of asymmetrical walking patterns when the perturbation is removed (i.e. symmetrical walking condition). This phenomenon, known as the after-effect, reflects that the central nervous system learns and retains new walking patterns suited to the asymmetric condition via predictive feedforward adaptation (Reisman *et al*., 2010). The after-effect eventually diminishes (i.e. de-adaptation) and persists longer with repeated adaptation paradigms (Reisman *et al*., 2013).

The neural mechanisms underlying split-belt locomotor adaptation have been extensively studied over the past two decades. Studies have shown that individuals with cerebellar damage (Morton & Bastian, 2006), following hemispherectomy (Choi *et al*., 2009), and after a stroke (Tyrell *et al*., 2014) exhibit slower adaptation or smaller after-effects than healthy young individuals. These findings suggest that the supraspinal structures are involved in split-belt locomotor adaptation (Hinton *et al*., 2020). Recently, part of the role of the cerebral cortex in split-belt locomotor adaptation was elucidated using high-density electroencephalography (EEG) (Jacobsen & Ferris, 2023). The EEG study revealed that the power around the beta band in multiple brain regions, such as the sensorimotor and posterior parietal cortices, temporarily decreased after exposure to asymmetric perturbation and after removal of the perturbation. As adaptation and de-adaptation progressed, the power levels in these brain regions returned to those observed during normal walking. These findings highlight the contribution of the supraspinal structure to the locomotor adaptation. In addition to changes in cortical activity, lower-limb muscle activity also showed temporary increases following exposure to asymmetric perturbation and removal of the perturbation (Ogawa *et al*., 2014). This increased muscle activity gradually decreases as adaptation and de-adaptation progress (Finley *et al*., 2013; Maclellan *et al*., 2014). Moreover, the activity of groups of muscles working together at certain times within one gait cycle (i.e., muscle synergy) has been shown to be modulated in split-belt locomotor adaptation (Oshima *et al*., 2022). In summary, both cortical and muscle activities changed with split-belt locomotor adaptation.

However, the functional coupling between cortical and muscle activities during split-belt locomotor adaptation remains unknown. Significant functional coupling (i.e. cortico- muscular coherence [CMC]) has been shown between cortical and lower-limb muscle activities (Petersen *et al*., 2012; Roeder *et al*., 2018; Spedden *et al*., 2019; Roeder *et al*., 2020; Yokoyama, Yoshida *et al*., 2020). Previous studies have reported that the CMC between the sensorimotor region and tibialis anterior (TA) muscle during steady-state walking shows gait phase-dependent modulation and is mainly observed in the alpha (8–12 Hz) and beta (12–32 Hz) bands (Roeder *et al*., 2018; Spedden *et al*., 2019; Roeder *et al*., 2020; Yokoyama, Yoshida *et al*., 2020). Given that CMC reflects communication between the cortex and muscles during movement (Baker, 2007; Liu *et al*., 2019; Bourguignon *et al*., 2019), CMC during walking may play a significant role in the neuromuscular control of walking. However, previous research has focused primarily on steady-state walking, leaving a gap in our understanding of CMC during split-belt locomotor adaptation. Therefore, the next step in further understanding of the neural mechanisms underlying split-belt locomotor adaptation will require a focus on the CMC.

We focused on the CMC between the sensorimotor cortex and the TA muscle because it is a key muscle for split-belt adaptation (Ogawa *et al*., 2014). CMC has been suggested to reflect both the descending drive from the cortex to the muscle and the sensory feedback from the muscle to the cortex (i.e. sensorimotor integration) (Baker, 2007; Liu *et al*., 2019; Bourguignon *et al*., 2019). A previous study showed that CMC decreases when the motor system is unable to accurately predict sensory feedback (Mendez-Balbuena *et al*., 2013). Based on this finding, we hypothesized that the CMC between the sensorimotor region and TA muscle would temporarily decrease after exposure to asymmetric perturbation and after removal of the perturbation because a mismatch between the predicted and actual sensory feedback was expected to occur. On the other hand, CMC was expected to increase as adaptation and de-adaptation progressed because the motor system would correct the mismatch and accurately predict sensory feedback. Results obtained in the present study reveal the cortical control of muscles during split-belt locomotor adaptation, deepening our knowledge of the neural control of gait and providing valuable insights for optimising gait rehabilitation.

## Methods

### Ethical approval

All study procedures were approved by the Ethics Committee of the University of Tokyo (approval number: 701-4) and conducted in accordance with the Declaration of Helsinki. The participants were comprehensively informed about the experimental design and any potential risks and provided written informed consent prior to their participation.

### Participants

Nineteen healthy young men (aged 23.6 ± 2.3 years, range: 21–30 years; height: 172.5 ± 6.8 cm; weight: 65.8 ± 11.2 kg) participated in this study. None of the participants had any neurological impairments or prior experience with split-belt locomotor adaptation.

### Experimental design

Figure 1A shows the walking paradigm using a split-belt treadmill (Bertec, USA). The paradigm consisted of three distinct periods: baseline (both belts moving at 0.6, 0.9, and 1.2 m/s for 5 min each), adaptation (right and left belts moving at 0.6 and 1.2 m/s, respectively, for 15 min), and washout (both belts moving at 0.9 m/s for 10 min). The right leg was designated as the “slow leg”, and the left leg as the “fast leg”. A 5-min seated rest was provided between the baseline and adaptation periods to reduce fatigue. The belts were temporarily stopped when the periods switched, and the acceleration and deceleration of the belts were set to 0.8 m/s^2^. The experimenter verbally informed the participants of the upcoming belt speed changes a few seconds before the changes. To minimise EEG signal contamination, the participants were instructed to look straight ahead without looking down as much as possible. After the walking paradigm, the participants remained seated for 2 min to record the resting EEG signals.

**Figure 1.**
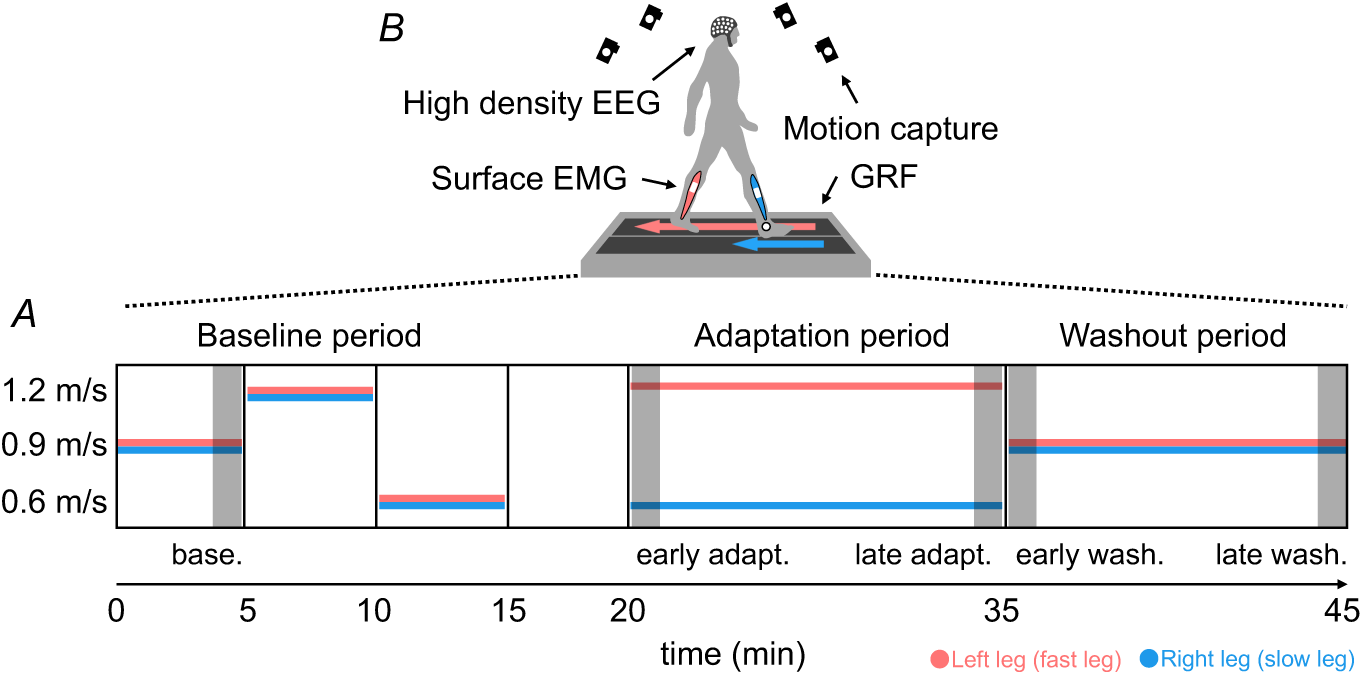
Experimental design. The horizontal axis represents time (A). The horizontal blue and red lines correspond to the slow leg and fast leg, respectively. The walking paradigm consisted of three periods: baseline, adaptation, and washout. During the baseline and washout periods, the belt speeds for both the right and left sides were identical. In the adaptation period, the belt speed ratio was 1:2 (0.6 m/s for the slow leg vs. 1.2 m/s for the fast leg). Each rectangle filled with dark grey represents sub-periods. Throughout the walking paradigm, high-density electroencephalogram (EEG), surface electromyogram (EMG) from the tibialis anterior muscle, ground reaction forces (GRF), and motion tracking data were recorded (B). base., baseline; early adapt., early adaptation; late adapt., late adaptation; early wash., early washout; late wash., late washout

### Data collection

Three-dimensional ground reaction force (GRF) data (mediolateral, anteroposterior, and vertical components) of each leg were recorded separately from the force plates mounted underneath each belt at a sampling rate of 1000 Hz using A/D conversion instruments (PowerLab 16/35, AD Instruments Inc., Australia) and recording software (LabChart, AD Instruments Inc., Australia) (Fig. 1B). The GRF data were used to identify heel contact (HC) and toe-off (TO) events while walking.

Motion-tracking data were collected at a sampling rate of 100 Hz using a motion-capture system (OptiTrack Motive, Natural Point Inc., USA) with 10 cameras (OptiTrack Flex3, Natural Point Inc., USA). Infrared reflective markers were attached to the lateral malleolus of each ankle, and motion-tracking data were used to compute the step length of each leg.

High-density EEG signals were recorded at a sampling rate of 500 Hz using an EEG system (actiChamp Plus, Brain Products, Germany), a cap (EasyCap, EASYCAP, Germany) with 160 active electrodes (actiCAP slim active electrodes, Brain Products, Germany), and recording software (BrainVision Recorder, Brain Products, Germany). The reference and ground electrodes were positioned at Fz and forehead, respectively. The impedance of most electrodes was maintained below 30 kΩ throughout the experiment.

Surface EMG signals were collected at a sampling rate of 2000 Hz from the muscle belly of the TA muscle of each leg using wireless active EMG electrodes (Pico, cometa, Italy) via a Wave Plus 16 channels amplifier (cometa, Italy) and recording software (EMG and Motion Tools, cometa, Italy). The skin on the muscle was cleaned with alcohol swabs to reduce impedance, and each electrode was secured with tape.

All data were synchronised using trigger signals delivered from the EMG recording system to the A/D conversion instruments (PowerLab) and from the A/D conversion instruments (PowerLab) to the EEG and motion capture systems before and after the walking paradigm. The data were then interpolated and aligned offline based on trigger timings (Artoni *et al*., 2017).

### Data analysis

Two participants were excluded from the analysis because of poor EMG signal quality caused by technical issues in one participant and the absence of adaptive changes in step length during the adaptation period in another participant. Data from the remaining participants were analysed using MATLAB R2021b (MathWorks Inc., USA).

### Ground reaction forces and motion-tracking data analysis

The GRF and motion-tracking data were smoothed using a low-pass filter with cutoff frequencies of 15 Hz (3rd-order Butterworth filter) and 6 Hz (4th-order Butterworth filter). The pre-processed GRF data were down-sampled to match the sampling rates of the other datasets. HC and TO events were identified based on the vertical component of the GRF, with the threshold set at 8% of each participant’s body weight (Sato & Choi, 2019).

Step-length symmetry (SLS) was computed from the identified HC events and motion-tracking data to quantify the degree of adaptation and washout. SLS was calculated from two consecutive steps and defined as the normalised difference between the step lengths of each leg using the following equation:

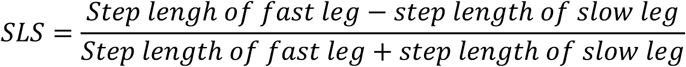

In this equation, the step length was the anterior-posterior distance between the ankle markers of each leg at the HC of the leading leg (Reisman et al., 2005). For example, the step length of the slow leg was calculated at the HC of the slow leg. A value of zero indicated that the step lengths of both legs were equal (i.e. symmetry), a negative value indicated that the step length of the slow leg was longer than that of the fast leg, and a positive value indicated that the step length of the fast leg was longer than that of the slow leg.

### EEG pre-processing

An overview of the EEG pre-processing steps is shown in Figure 2A. The EEG analysis was conducted using custom scripts incorporating EEGLAB v2022.0 functions. The recorded EEG signals were high-pass filtered at 1 Hz and low-pass filtered at 200 Hz using the “eegfiltnew” function (Yokoyama, Kaneko *et al*., 2020). The filtered EEG data were resampled at 256 Hz. Noisy channels were removed based on the following criteria (Gwin *et al*., 2011; Yokoyama, Kaneko *et al*., 2020): (1) a standard deviation (SD) exceeding 1000 μV, (2) kurtosis greater than five SD from the mean. Moreover, artefact subspace reconstruction was conducted using “asr_calibrate” and “asr_process” functions to eliminate high-amplitude artefacts from the EEG data during the walking paradigm. EEG data recorded in a seated position were used as the baseline, and the cutoff criterion was set at SD = 20 (Gorjan *et al*., 2022). The EEG data were subsequently re-referenced to a common average reference (Roeder *et al*., 2018; Yokoyama, Kaneko *et al*., 2020). AC power noise (50 and 100 Hz) was attenuated using the “cleanline” function. The EEG data were divided into 3.0 s epochs surrounding the HC events for each leg, spanning from 1 s before to 2 s after the HC events. Outlier epochs were excluded based on joint probability (6 SD for single channels and 2 SD for all channels) using the “pop_jointprob” function. Following pre-processing, the EEG data were decomposed using the CUDAICA implementation of the Infomax independent component analysis (ICA) to identify the brain and non-brain sources mixing in the EEG data (“pop_runica” function). EEG data from all periods were concatenated for ICA. The resulting independent components (ICs) were automatically classified and labelled using the “pop_iclabel” function. ICs labelled as “brain” and “other” and the remaining ICs showing < 75% probability (Studnicki, Seidler *et al*., 2023) were retained for further analysis. The remaining ICs were projected back onto the channels. This procedure cleaned the EEG signals from eye blinks, muscle noise, heart artefacts, and other contaminants, while preserving brain-related activity as much as possible. Finally, the Hjorth Laplacian derivation was applied to the EEG data of each electrode, using the mean value calculated from the electrodes surrounding the target electrode as a reference. (Kanazawa *et al*., 2014; Yokoyama, Yoshida *et al*., 2020).

**Figure 2.**
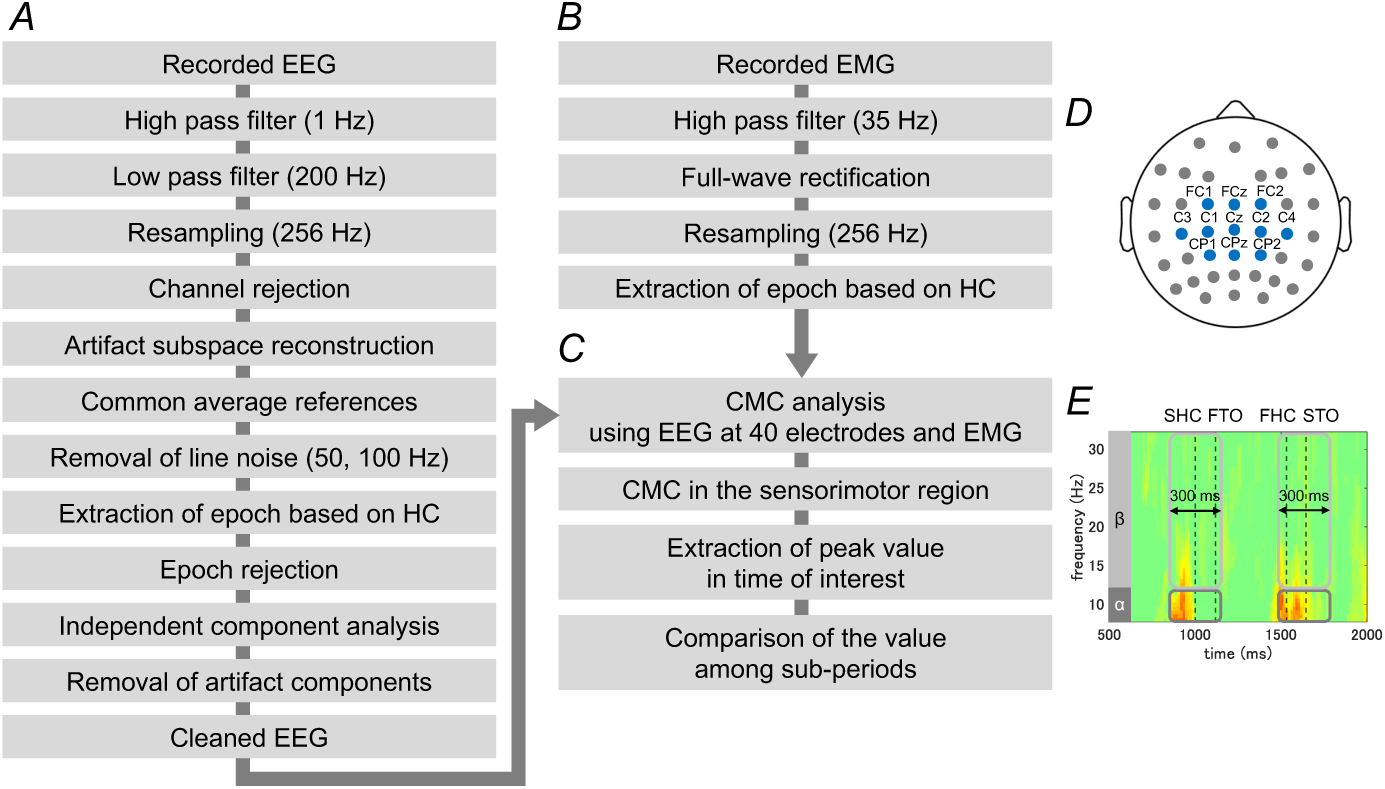
Overview of cortico-muscular coherence analysis. Flowchart of electroencephalogram (EEG) (A) and electromyogram (EMG) (B) pre-processing for cortico-muscular coherence (CMC) analysis (C). The electrodes used to calculate CMC are shown in (D). Electrodes in the sensorimotor region are shown in blue. Each rectangle illustrates the time of interest within a single gait cycle (E). This panel shows the time of interest for the slow leg. In this figure, the horizontal axis represents time (ms), and the vertical axis represents frequency (Hz). The alpha band ranged 8–12 Hz, and the beta band ranged 12–32 Hz. Dashed vertical lines indicate heel contact and toe-off events for each leg. SHC, slow leg heel contact; FTO, fast leg toe-off; FHC, fast leg heel contact; STO, slow leg toe-off

### Cortico-muscular coherence analysis

The recorded EMG signals were high-pass filtered at 35 Hz (4th-order Butterworth filter) to reduce motion artefacts (Shuman *et al*., 2017) and then full-wave rectified (Fig. 2B) (Yokoyama, Yoshida *et al*., 2020). The rectified EMG data were resampled to 256 Hz and analysed along with the cleaned EEG data. The “newcrossf” function was used to compute the event-related coherence between EEG and EMG data (Studnicki & Ferris, 2023) (Fig. 2C). Initially, the CMC was calculated between the EEG data from 40 electrodes positioned across the entire scalp (Fig. 2D) and the TA EMG data from each leg to determine the scalp distribution of the CMC. Because a high CMC was observed between the EEG data from the electrodes near the sensorimotor region and the TA EMG data, subsequent analyses focused on 11 electrodes near the sensorimotor region (points highlighted in blue in Fig. 2D) (Jacobs *et al*., 2015; Yokoyama *et al*., 2019). Significances in the individual event-related coherence at each electrode were tested using a bootstrapping method (α = 0.05, shuffled in the time dimension, 2000 iterations) implemented in the “newcrossf” function. Next, the CMC between the sensorimotor region and the TA muscle during walking has been primarily observed in the alpha (8–12 Hz) and beta (12–32 Hz) bands in two gait phases (around HC and TO events) (Roeder *et al*., 2018, 2020; Spedden *et al*., 2019; Yokoyama, Yoshida *et al*., 2020); therefore, four time of interest were set as follows: alpha and beta bands around HC and TO events (Fig. 2E). The duration of each time of interest was 300 ms, centred on HC or TO events. The maximal CMC value at each time of interest was extracted from individual event-related coherence (Gwin & Ferris, 2012; Ushiyama *et al*., 2017; Yokoyama, Yoshida *et al*., 2020). The maximal value among the 11 electrodes at each time of interest was used for statistical analyses. The maximum value was normalised using an arc hyperbolic tangent transformation (Halliday *et al*., 1995; Ushiyama *et al*., 2010, 2017).

### Statistical analysis

The baseline, adaptation, and washout periods were segmented into the following five sub-periods (each rectangle filled with dark grey in Fig. 1A): baseline, early adaptation, late adaptation, early washout, and late washout, to investigate changes in SLS and CMC during split-belt locomotor adaptation. Each sub-period was segmented using data from all gait cycles for SLS. In contrast, each sub-period was segmented based on the EEG data after pre-processing (i.e. some gait cycles were removed owing to epoch rejection) for the CMC. In both cases, each sub-period comprised 70 gait cycles. Participants who did not show significant CMC in any sub-period were excluded from the statistical analysis (Gennaro & de Bruin, 2020). The number of participants for each statistical analysis is presented in Tables 1 and 2. The normality of the data was evaluated using the Shapiro-Wilk test. The Fisher-Pitman permutation test of matched pairs with 50000 permutations was used because a normal distribution was not observed in most cases. The *p*-values were corrected for multiple comparisons using false discovery rate (FDR) correction (Benjamini & Hochberg, 1995). The significance level was set at *a* = 0.05. The effect size on the differences between the sub-periods was indicated using Cliff’s delta. The effect size criteria were quantified with values of 0.11, 0.28, and 0.43 indicating small, medium, and large effects, respectively (Vargha & Delaney, 2000).

**Table 1.** Statistical information for Figure 3. Significant *p* values are indicated in bold italics.

| Corrected $p$ value (Fisher-Pitman permutation test of matched pairs with FDR correction) and effect size (Cliff's delta) | | | | | | | | | | |
| --- | --- | --- | --- | --- | --- | --- | --- | --- | --- | --- |
| SLS | n = 17 | sub-period | early adaptation |  | late adaptation |  | early washout |  | late washout |  |
|  |  | baseline | < ***0.001*** | 1.000 | < ***0.001*** | 0.721 | < ***0.001*** | 0.978 | < ***0.001*** | 0.493 |
|  |  | early adaptation |  |  | < ***0.001*** | 0.971 | < ***0.001*** | 1.000 | < ***0.001*** | 1.000 |
|  |  | late adaptation |  |  |  |  | < ***0.001*** | 0.971 | < ***0.001*** | 0.801 |
|  |  | early washout |  |  |  |  |  |  | < ***0.001*** | 0.904 |

**Table 2.**
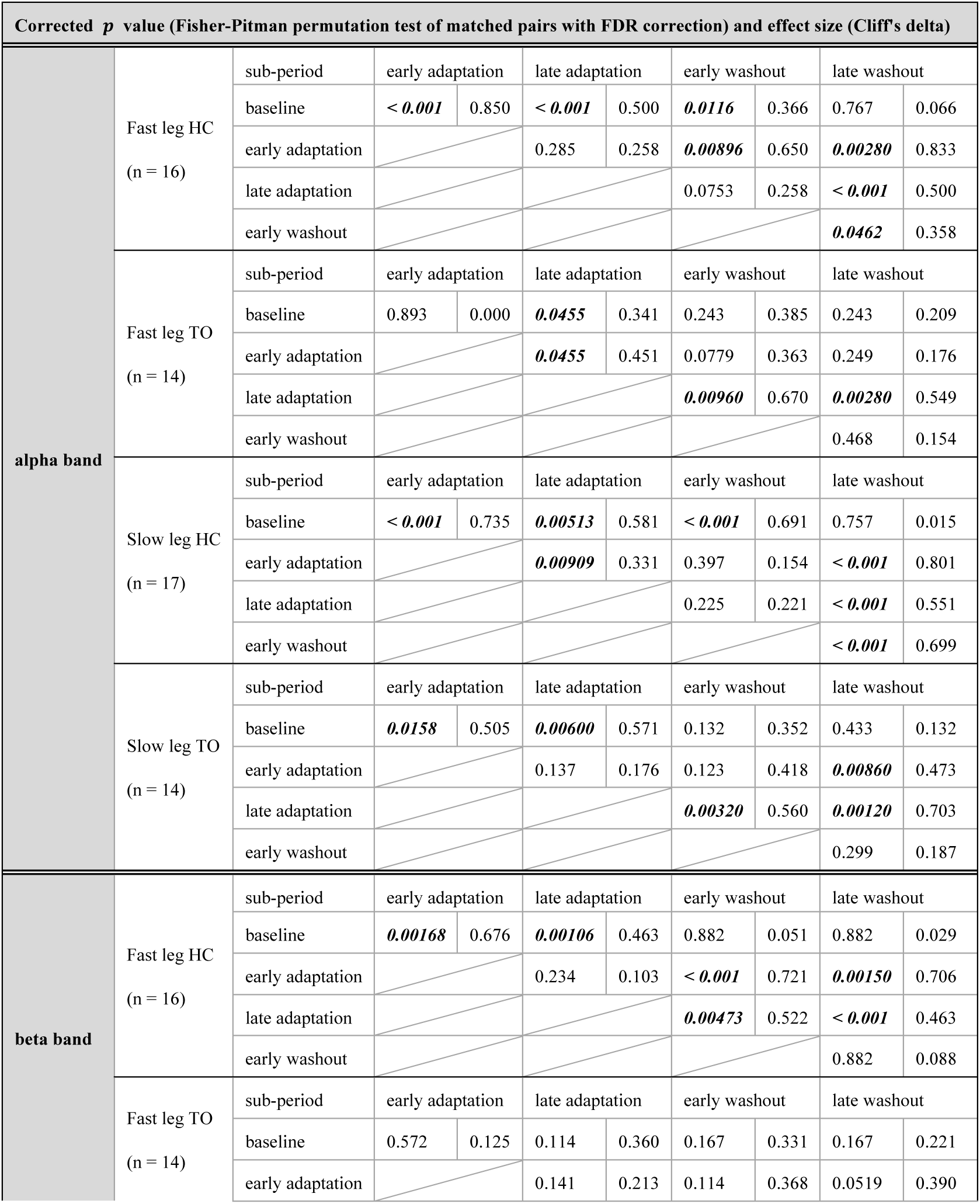
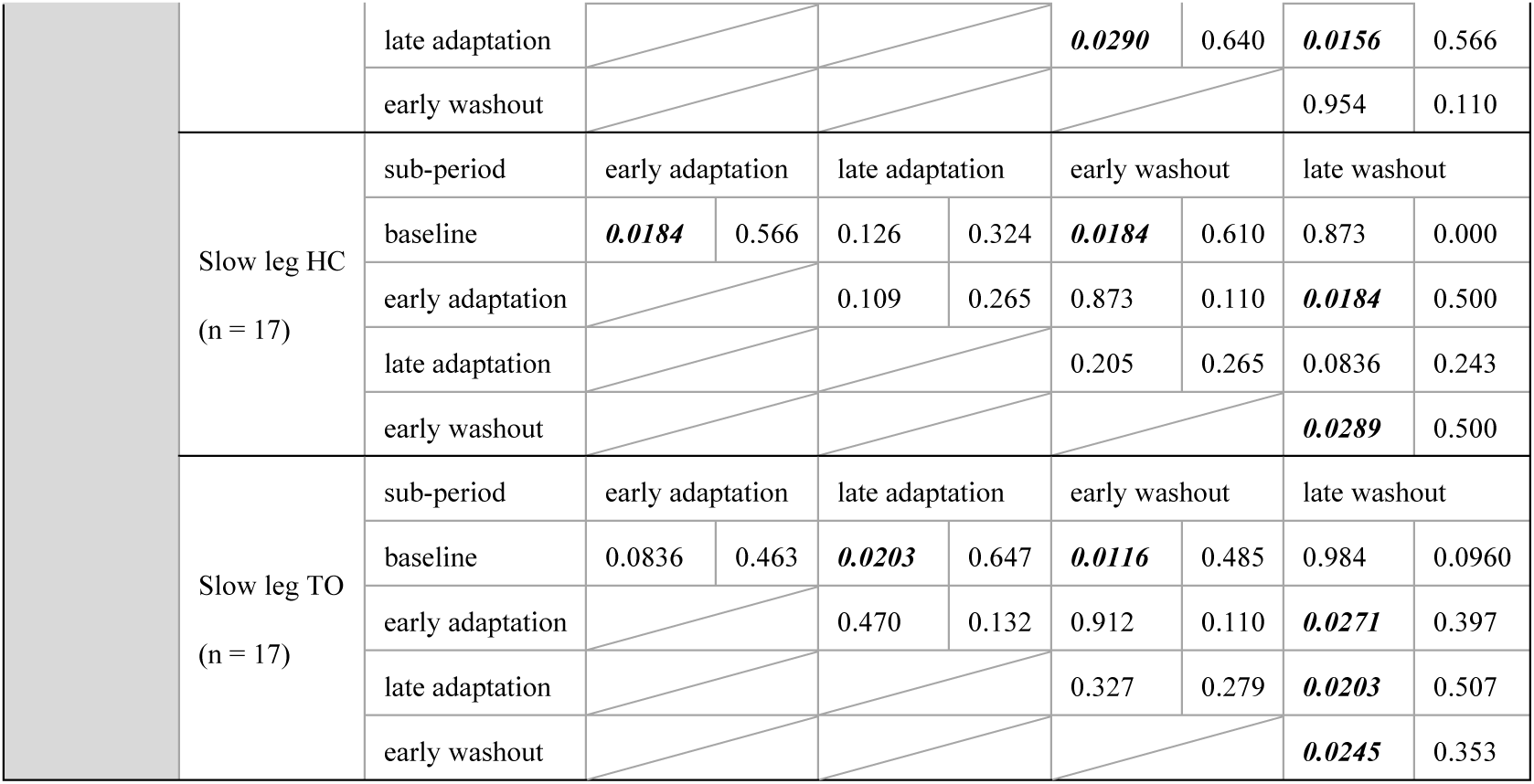
Statistical information for Figure 5. Significant *p* values are indicated in bold italics.

## Results

### Step length symmetry

The adaptive changes in the SLS observed in the walking paradigm were as follows: SLS was almost zero at baseline (Fig. 3A). Immediately after exposure to the adaptation period, it showed a large negative value, gradually returning to zero throughout the adaptation period. In contrast, SLS showed a positive value in the early washout and then converged toward zero in the late phase of the washout period. The statistical data are summarised in Table 1. The SLS at baseline was significantly higher than that at early adaptation (*p*_adj_ < 0.001) and late adaptation (*p*_adj_ < 0.001), and significantly lower than that at early washout (*p*_adj_ < 0.001) and late washout (*p*_adj_ < 0.001) (Fig. 3B). The SLS during early adaptation was significantly lower than that during late adaptation (*p*_adj_ < 0.001), early washout (*p*_adj_ < 0.001), and late washout (*p*_adj_ < 0.001). Additionally, the SLS in the late adaptation was significantly lower than those in the early washout (*p*_adj_ < 0.001) and late washout (*p*_adj_ < 0.001). Finally, the SLS in early washout was significantly higher than that in late washout (*p*_adj_ < 0.001).

**Figure 3.**
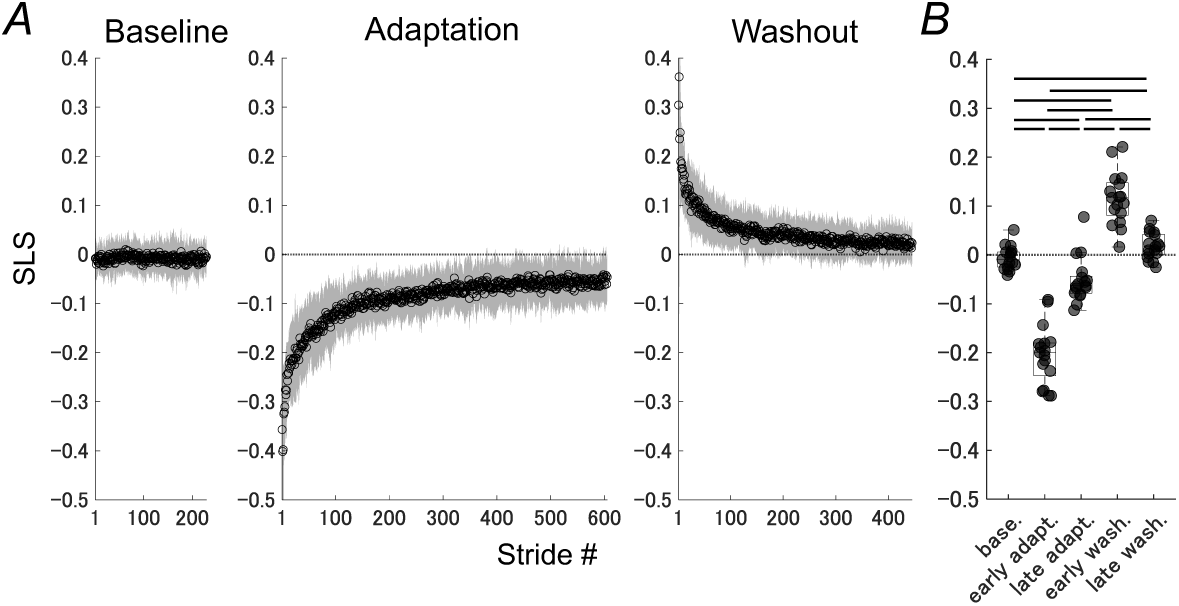
Step length symmetry. The vertical axis represents step length symmetry (SLS), and the horizontal axis shows the number of strides (A) and sub-periods (B). A value of zero on the vertical axis (dotted lines) indicates that the step lengths of the slow and fast legs were equal (i.e. symmetry). Each circle indicates the SLS for each stride, and the grey-shaded areas surrounding each circle indicate the standard deviation (A). The line within each box denotes the median value, and the ends of each box represent the 25th and 75th percentiles (B). Each circle represents the data from each participant. Horizontal bars indicate significant differences between the sub-periods (*p* < 0.05). Statistical information is presented in Table 1. base., baseline; early adapt., early adaptation; late adapt., late adaptation; early wash., early washout; late wash., late washout

### Scalp topography and time-frequency plot of cortico-muscular coherence

The ground-averaged scalp topography of the CMC is shown in Fig 4A–H. This topography was obtained using the CMC calculated from 40 electrodes distributed across the entire scalp. High CMC was predominantly observed near the sensorimotor region (highlighted by grey circles). The CMC in this region changed with split-belt locomotor adaptation.

**Figure 4.**
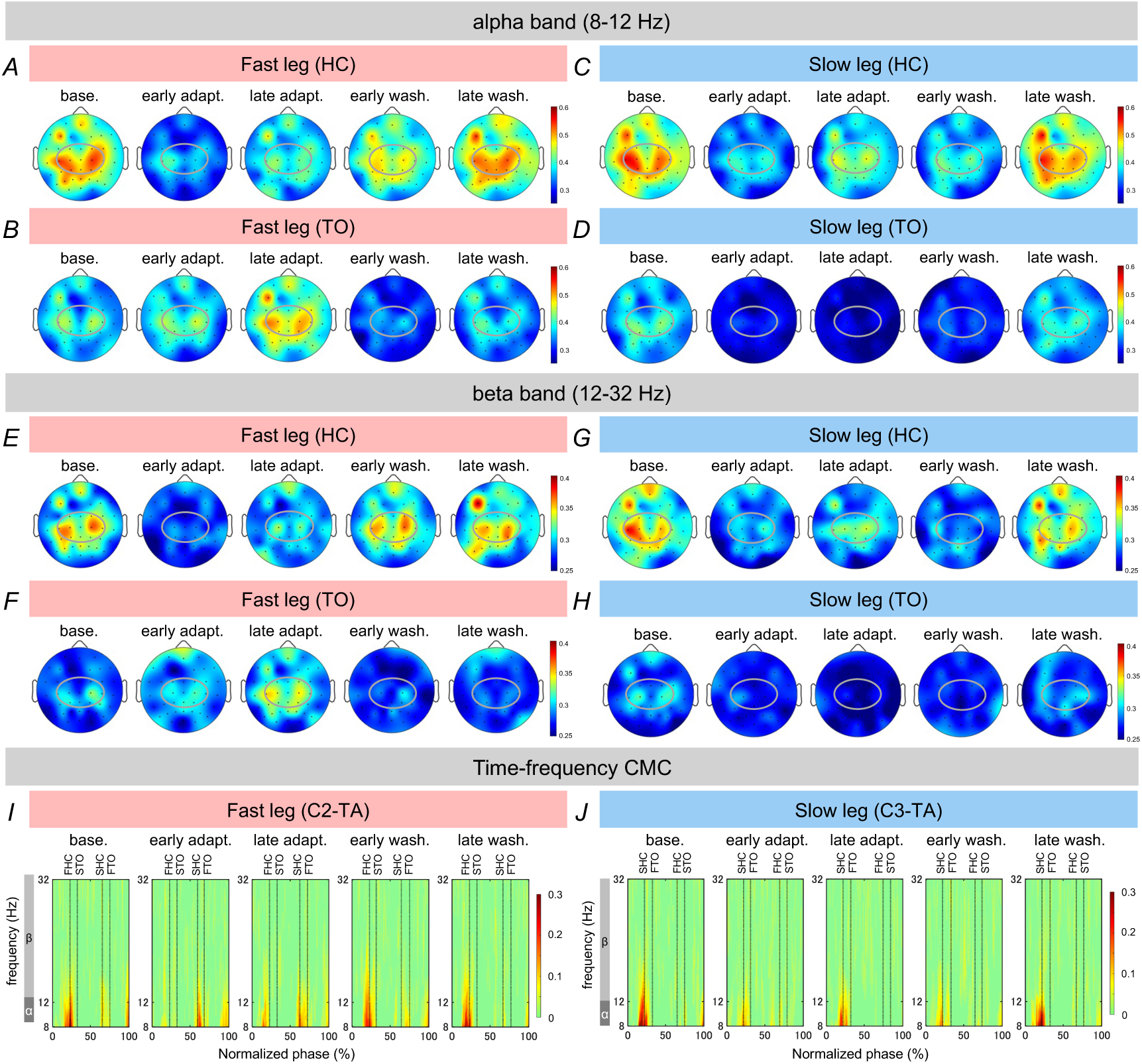
Scalp topography and time-frequency plot of cortico-muscular coherence. The upper (A–D) and middle panels (E–H) represent the topography of cortico-muscular coherence (CMC) in the alpha (8–12 Hz) and beta (12–32 Hz) bands, respectively. The panels on the left (A, B, E, and F) and right (C, D, G, and H) show scalp topography of the fast and slow legs, respectively. The lower panels (I and J) show the time-frequency plots of CMC for the fast and slow legs, respectively. The vertical axis represents the frequency (Hz), and the horizontal axis represents the normalised time. The dashed vertical lines within the plot indicate the heel contact and toe-off events for each leg. The alpha band ranged 8–12 Hz, and the beta band ranged 12–32 Hz. Time-frequency plots were created using the most frequently selected electrode across the participants when extracting the maximal CMC values. C2 was used for the fast and C3 for the slow leg, respectively. base., baseline; early adapt., early adaptation; late adapt., late adaptation; early wash., early washout; late wash., late washout; HC, heel contact; TO, toe-off; SHC, slow leg heel contact; FTO, fast leg toe-off; FHC, fast leg heel contact; STO, slow leg toe-off

Specifically, except for alpha CMC in fast leg TO (Fig. 4B), alpha CMC was reduced in early adaptation, late adaptation, and early washout compared with baseline and late washout (Fig. 4A, C, and D). In contrast, alpha CMC in fast leg TO increased in the late adaptation compared to the other sub-periods (Fig. 4B). In the late washout, the alpha CMC returned to levels similar to baseline (Fig. 4A–D). Although the magnitude of CMC was generally smaller in the beta band than in the alpha band (Fig. 4E–H), the overall trends in CMC during split-belt locomotor adaptation were consistent across both bands.

Figure 4 also presents the time-frequency plot of the CMC. The time-frequency plot in the fast and slow legs was calculated using EEG signals at C2 and C3 and EMG signals from the TA muscle (Fig. 4I and J). The CMC in each leg was high in the alpha (8–12 Hz) and low- beta (approximately 12–20 Hz) bands, particularly around the HC and TO.

### Comparison of cortico-muscular coherence among sub-periods

Figure 5 shows the CMC values at each sub-period during the walking paradigm, and Table 2 summarises the corresponding statistical results. Based on the results shown in Figure 5 and Table 2, we report the results focusing on the following three points that are mainly related to the evaluation of the locomotor adaptation process: 1) differences from baseline, 2) changes during the adaptation period, and 3) changes during the washout period.

**Figure 5.**
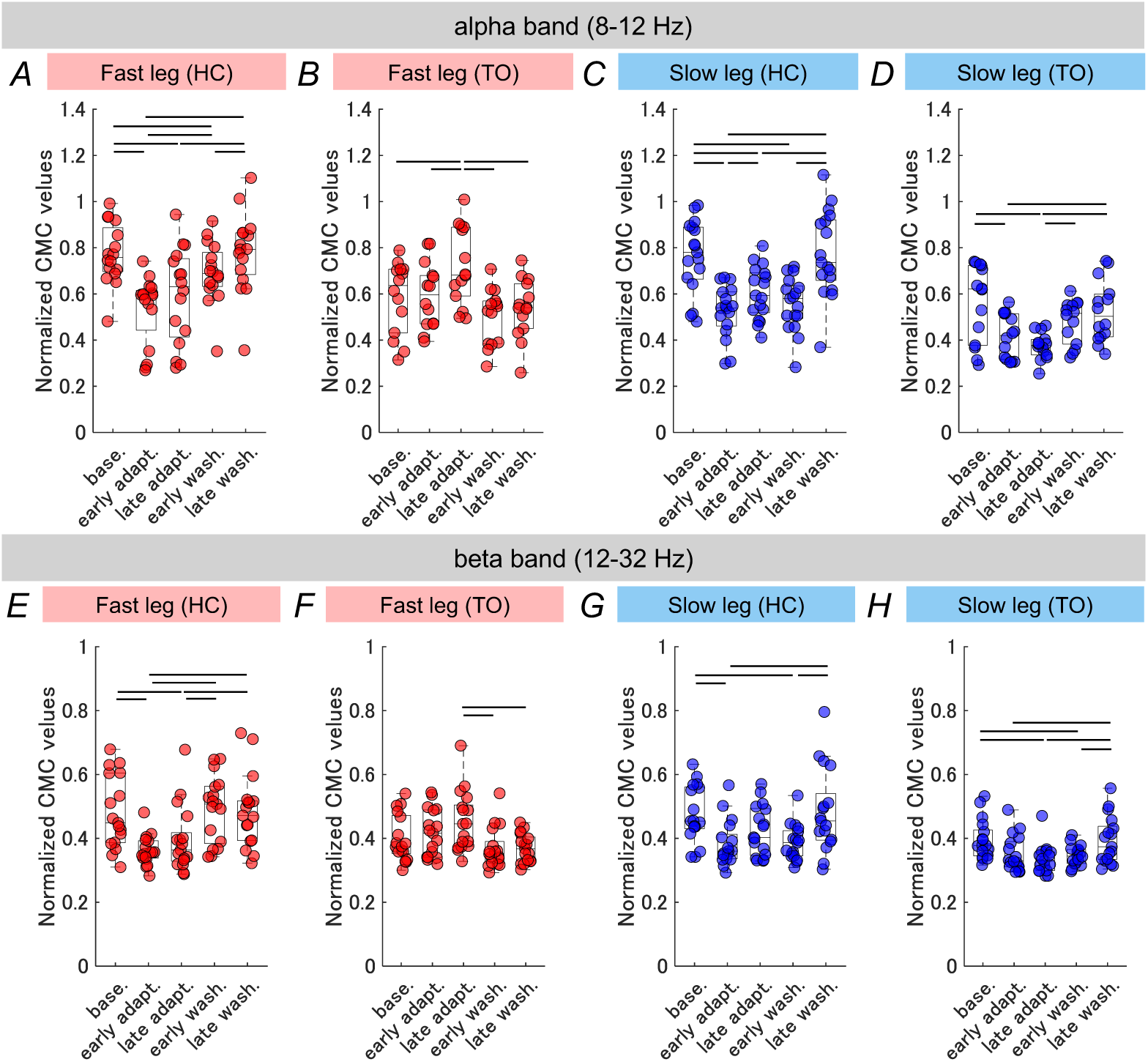
Comparison of cortico-muscular coherence among sub-periods. The upper panels (A–D) and lower panels (E–H) display cortico-muscular coherence (CMC) in the alpha (8–12 Hz) and beta (12–32 Hz) bands, respectively. The left (A, B, E, and F) and right panels (C, D, G, and H) represent data from the fast and slow legs, respectively. The vertical axis represents the normalised CMC value, and the horizontal axis represents the sub-periods. The line within each box represents the median value, and the ends of each box represent the 25th and 75th percentiles. Each circle represents the data from each participant. Significant differences between the sub-periods are indicated by horizontal bars (*p* < 0.05). The statistical information is summarised in Table 2. base., baseline; early adapt., early adaptation; late adapt., late adaptation; early wash., early washout; late wash., late washout; HC, heel contact; TO, toe-off

1) Differences from the baseline

The alpha CMC in the early adaptation and late adaptation was significantly lower than at baseline in the fast leg HC, slow leg HC, and slow leg TO (fast leg HC base. vs. early adapt.: *p*_adj_ < 0.001; fast leg HC base. vs. late adapt.: *p*_adj_ < 0.001; slow leg HC base. vs. early adapt.: *p*_adj_ < 0.001; slow leg HC base. vs. late adapt.: *p*_adj_ = 0.00513; slow leg TO base. vs. early adapt.: *p*_adj_ = 0.0158; slow leg TO base. vs. late adapt.: *p*_adj_ = 0.00600) (Fig. 5A, C, and D). For the fast leg TO, the alpha CMC in the late adaptation was significantly higher than that at baseline (*p*_adj_ = 0.0455) (Fig. 5 B). In the early washout, the alpha CMC in the fast leg HC and slow leg HC was significantly lower than that at baseline (fast leg HC: *p*_adj_ = 0.0116; slow leg HC: *p*_adj_ < 0.001) (Fig. 5A and C). In contrast, no significant differences were observed between baseline and late washout in each leg (Fig. 5A-D).

The beta CMC in early adaptation was significantly lower than that at baseline in the fast leg and slow leg HC (fast leg HC: *p*_adj_ = 0.00168; slow leg HC: *p*_adj_ = 0.0184) (Fig. 5E and G). The beta CMC during late adaptation was also significantly lower than that at baseline in the fast leg HC and slow leg TO (fast leg HC: *p*_adj_ = 0.00106; slow leg TO: *p*_adj_ = 0.0203) (Fig. 5E and H). The beta CMC in the early washout was significantly lower than that in the baseline in the slow leg HC and TO (slow leg HC: *p*_adj_ = 0.0184; slow leg TO: *p*_adj_ = 0.0116) (Fig. 5G and H). Similar to alpha CMC, statistical analysis did not show significant differences between baseline and late washout (Fig. 5E-H).

2) Changes during the adaptation period

Significant increases from early adaptation to late adaptation were observed for alpha CMC in the fast leg TO and slow leg HC (fast leg TO: *p*_adj_ = 0.0455; slow leg HC: *p*_adj_ = 0.00909) (Fig. 5B and C).

3) Changes during the washout period

Alpha CMC in the fast and slow leg HC significantly increased (fast leg HC: *p*_adj_ = 0.0462; slow leg HC: *p*_adj_ < 0.001) (Fig. 5A and C). In addition, significant increases in beta CMC were observed in the slow leg HC and TO (slow leg HC: *p*_adj_ = 0.0289; slow leg TO: *p*_adj_ = 0.0245) (Fig. 5G and H).

## Discussion

We investigated the CMC between the sensorimotor region and TA muscle in split-belt locomotor adaptation. Our results for the SLS (Fig. 3) and CMC (Fig. 4) during steady-state walking (baseline) are consistent with those reported in previous studies (Reisman *et al*., 2005; Choi *et al*., 2009; Roeder *et al*., 2018, 2020; Spedden *et al*., 2019; Yokoyama, Yoshida *et al*., 2020). The magnitude of the CMC in the alpha and beta bands was greater in the sensorimotor region and was enhanced around the HC and TO events (Fig. 4), supporting the validity of our results.

We hypothesised that the CMC would temporarily decrease after exposure to asymmetric perturbation and after removal of the perturbation. Our results partially supported this hypothesis. Specifically, alpha CMC in fast leg HC, slow leg HC, and slow leg TO (Fig. 5A, C, D) and beta CMC in fast leg HC and slow leg HC (Fig. 5 E, G) in the early adaptation were significantly lower than those at baseline. Additionally, alpha CMC in fast leg HC and slow leg HC (Fig. 5A, C) and beta CMC in slow leg HC and TO (Fig. 5 G, H) during the early washout were significantly lower than those at baseline. When external conditions are predictable (i.e. baseline), the motor system can utilise sensorimotor information from previous experiences to generate appropriate motor commands, leading to efficient sensorimotor information processing between the cortex and muscles (Baker, 2007; Liu *et al*., 2019; Bourguignon *et al*., 2019).

However, the original and newly learned walking patterns were disturbed in early adaptation and early washout owing to the application of the asymmetrical perturbation and the subsequent removal of this perturbation. This results in a mismatch between the predicted and actual sensory feedback. The CMC has been reported to be reduced when such a mismatch occurs in a force control task using the upper limbs (Mendez-Balbuena *et al*., 2013), supporting our interpretation of decreased CMC.

The observed decrease in CMC during early adaptation and early washout could be attributed to cortical modulation, which is supported by the following findings: A recent EEG study using a split-belt paradigm demonstrated reductions in power in multiple brain regions, including the sensorimotor cortex, during early adaptation and early washout compared to baseline (Jacobsen & Ferris, 2023). Such cortical desynchronisation is associated with reduced CMC (Kristeva *et al*., 2007). On the other hand, a reduction in cortico-muscular coupling may not negatively impact the motor system. Although CMC is useful for maintaining stable motor output (Kristeva *et al*., 2007), the release of tight coupling between oscillatory activities in the motor system may allow for more flexible motor control (Halliday *et al*., 2003; Baker, 2007; Hug *et al*., 2021). Thus, our findings suggest that switching the cortico-muscular coupling state is a neural strategy adopted by the motor system to cope with asymmetrical perturbation and maintain stable walking.

Previous studies on split-belt locomotor adaptation have consistently shown that adaptive measures gradually change and reach a steady state as adaptation progresses, followed by the disappearance of after-effects with de-adaptation. (Reisman *et al*., 2005; Choi & Bastian, 2007; Finley *et al*., 2013; Ogawa *et al*., 2014; Maclellan *et al*., 2014; Oshima *et al*., 2022; Jacobsen & Ferris, 2023; Refy *et al*., 2023). Our second hypothesis was that CMC would increase with adaptation and de-adaptation, which was partially supported by the present results. During the adaptation period, the alpha CMC significantly increased in fast leg TO and slow leg HC (Fig. 5B, C). During the washout period, alpha CMC in fast leg HC and slow leg HC (Fig. 5A, C), and beta CMC in the slow leg HC and TO significantly increased (Fig. 5G, H).

First, we consider the relationship between increased CMC and the adaptation and de-adaptation processes. A previous study showed that the CMC between the sensorimotor region and a hand muscle increased with the learning of a visuomotor task (Mendez-Balbuena *et al*., 2012), which was similar to our results. As adaptation progresses, the motor system likely corrects the mismatch between the predicted and actual sensory feedback through trial-and-error practice, forming a new motor program suited to walking under asymmetric perturbation (Reisman *et al*., 2010). During the washout period, the motor system would recall a motor program appropriate for a normal walking environment with de-adaptation. Thus, the increase in CMC during the adaptation and de-adaptation periods may have resulted from the establishment and re-establishment of efficient sensorimotor integration, accompanied by the construction of new motor programs and the recall of existing motor programs (Perez *et al*., 2006; Baker, 2007; Ushiyama *et al*., 2017). However, it is important to note that not all gait phases exhibited significant changes in CMC with adaptation and de-adaptation. During the adaptation period, alpha CMC in fast leg HC and slow leg TO (Fig. 5A, D) and beta CMC in fast leg HC and TO and slow leg HC and TO (Fig. 5E–H) did not show a significant increase. Similarly, during the washout period, alpha CMC in fast leg TO and slow leg TO (Fig. 5B, D), and beta CMC in the fast leg HC and TO (Fig. 5E, F) did not increase significantly. These results suggest that sensorimotor integration may not have been updated or required in these specific gait phases. Alternatively, the motor system may require more time to establish efficient sensorimotor integration during these gait phases. Thus, the differences in the updating of sensorimotor integration depending on the leg, frequency bands, and gait phases might be one of the characteristics of split-belt locomotor adaptation.

Second, we expand the discussion of the increase in CMC observed during the adaptation period. The gait phases showing an increase in CMC were the slow leg HC and fast leg TO (Fig. 5B, C), corresponding to the fast leg double-support phases (i.e. the time from slow leg HC to fast leg TO). This suggests that adaptation to asymmetric perturbation is characterised by modulation of the CMC in the fast leg double support phases. There is a plausible explanation for the increased CMC in slow leg HC (Fig. 5C). Previous studies have indicated that HC control is critical not only for stabilising ankle joints and maintaining stable walking (Christensen *et al*., 2000) but also for split-belt locomotor adaptation (Malone *et al*., 2012). Additionally, the slow leg likely serves as a “reference leg” in split-belt adaptation, which is supported by studies detailing the kinetics and after-effects of split-belt locomotor adaptation (Vasudevan & Bastian, 2010; Ogawa *et al*., 2014; Hamzey *et al*., 2016). Vertical force has been shown to be larger on the slow leg than on the fast leg during adaptation (Ogawa *et al*., 2014). In addition, the anterior braking force at the slow leg HC increased rapidly during the initial adaptation period (Ogawa *et al*., 2014), which disrupted smooth walking movement. Moreover, studies examining the effects of walking speed on the magnitude of after-effects found that the after-effect was most pronounced when the slower speed used during the adaptation period was reapplied, compared to other speeds (Vasudevan & Bastian, 2010; Hamzey *et al*., 2016). This may be attributed to the fact that the stance duration of the slow leg is longer than that of the fast leg under asymmetric perturbation, causing the motor system to focus more on the slow leg and gather more sensory information from it (Vasudevan & Bastian, 2010). Therefore, given the importance of slow leg HC in adaptation to asymmetric perturbation, the motor system might intensively establish sensorimotor integration in the gait phase, resulting in increased CMC.

On the other hand, the magnitude of CMC in fast leg TO was significantly greater in the late adaptation than not only in early adaptation but also in baseline (Fig. 5B). One possible interpretation of these results is a change in muscle contraction patterns with adaptation to asymmetric perturbation. A previous study reported that the CMC is modulated depending on muscle contraction patterns (Riddle & Baker, 2006). Specifically, CMC increased when the muscle contraction patterns were steady. The slow leg moved slowly during the adaptation period and spent a long time in the stance phase, which may have led to a longer swing time for the fast leg. Consequently, the muscle contraction pattern in the fast leg likely became steady, contributing to an increased CMC during late adaptation.

We investigated the modulation of CMC between the sensorimotor area and the TA muscle, a key muscle in not only walking but also split-belt locomotor adaptation (Petersen *et al*., 2010; Ogawa *et al*., 2014). Our findings provide a foundation for further research on cortico-muscle coupling in the other lower-limbs during locomotor adaptation. Walking also relies on the coordinated activity of multiple muscles, which is known as muscle synergy (Ivanenko *et al*., 2004). Our group recently identified adaptive changes in muscle synergies during split-belt walking (Oshima *et al*., 2022) and also demonstrated the cortical correlates of locomotor muscle synergies during steady-state walking (Yokoyama *et al*., 2019). Integrating these insights into future studies may further our understanding of cortical control during locomotor adaptation.

## Conclusion

This study characterised the modulation of cortico-muscular coupling associated with split-belt locomotor adaptation. Our findings revealed that the CMC between the sensorimotor region and the TA muscle temporarily decreased depending on the leg, frequency bands, and gait phases after exposure to the asymmetric perturbation and removal of the perturbation. The CMC then increased depending on the leg, frequency bands, and gait phases as adaptation and de-adaptation to the perturbation progressed. These results suggest that flexible updating of cortico-muscular coupling in the motor system is a key mechanism underlying locomotor adaptation in humans. The present findings expand our understanding of locomotor adaptation and provide valuable insights into gait rehabilitation.

## Additional information

### Author contributions

The experiments in the present study were performed in the laboratories (rooms 104 and 116) in the University of Tokyo, Komaba Campus, Bldg. 9. AO, HY, and NK contributed to conception of this work. AO, HY, NK, and RT contributed to acquisition, analysis, and interpretation of data for this work. AO contributed to drafting this paper and HY, NK, RT, KT, and KN contributed to revising this paper critically. All authors approved the final version of the manuscript, and agreed to be accountable for all aspects of the work in ensuring that questions related to the accuracy or integrity of any part of the work are appropriately investigated and resolved. All persons designated as authors qualify for authorship, and all those who qualify for authorship are listed.

### Data availability statement

The raw data supporting the conclusions of this study are available from the corresponding author upon reasonable request.

## References

Artoni F, Barsotti A, Guanziroli E, Micera S, Landi A & Molteni F (2017). Effective synchronization of EEG and EMG for mobile brain/Body Imaging in clinical settings. Front Hum Neurosci 11, 652.

Baker SN (2007). Oscillatory interactions between sensorimotor cortex and the periphery. Curr Opin Neurobiol 17, 649–655.

Bastian AJ (2008). Understanding sensorimotor adaptation and learning for rehabilitation. Curr Opin Neurol 21, 628–633.

Benjamini Y & Hochberg Y (1995). Controlling the false discovery rate: A practical and powerful approach to multiple testing. J R Stat Soc 57, 289–300.

Bourguignon M, Jousmäki V, Dalal SS, Jerbi K & De Tiège X (2019). Coupling between human brain activity and body movements: Insights from non-invasive electromagnetic recordings. Neuroimage 203, 116177.

Choi JT & Bastian AJ (2007). Adaptation reveals independent control networks for human walking. Nat Neurosci 10, 1055–1062.

Choi JT, Vining EPG, Reisman DS & Bastian AJ (2009). Walking flexibility after hemispherectomy: split-belt treadmill adaptation and feedback control. Brain 132, 722– 733.

Christensen LO, Petersen N, Andersen JB, Sinkjaer T & Nielsen JB (2000). Evidence for transcortical reflex pathways in the lower limb of man. Prog Neurobiol 62, 251–272.

Finley JM, Bastian AJ & Gottschall JS (2013). Learning to be economical: the energy cost of walking tracks motor adaptation. J Physiol 591, 1081–1095.

Gennaro F & de Bruin ED (2020). A pilot study assessing reliability and age-related differences in corticomuscular and intramuscular coherence in ankle dorsiflexors during walking. Physiol Rep 8, e14378.

Gorjan D, Gramann K, De Pauw K & Marusic U (2022). Removal of movement-induced EEG artifacts: current state of the art and guidelines. J Neural Eng; DOI: 10.1088/1741-2552/ac542c.

Gwin JT & Ferris DP (2012). Beta- and gamma-range human lower limb corticomuscular coherence. Front Hum Neurosci 6, 258.

Gwin JT, Gramann K, Makeig S & Ferris DP (2011). Electrocortical activity is coupled to gait cycle phase during treadmill walking. Neuroimage 54, 1289–1296.

Halliday DM, Conway BA, Christensen LOD, Hansen NL, Petersen NP & Nielsen JB (2003). Functional coupling of motor units is modulated during walking in human subjects. J Neurophysiol 89, 960–968.

Halliday DM, Rosenberg JR, Amjad AM, Breeze P, Conway BA & Farmer SF (1995). A framework for the analysis of mixed time series/point process data—Theory and application to the study of physiological tremor, single motor unit discharges and electromyograms. Progress in Biophysics and Molecular Biology 64, 237–278. Available at: 10.1016/s0079-6107(96)00009-0.

Hamzey RJ, Kirk EM & Vasudevan EVL (2016). Gait speed influences aftereffect size following locomotor adaptation, but only in certain environments. Exp Brain Res 234, 1479–1490.

Hinton DC, Conradsson DM & Paquette C (2020). Understanding Human Neural Control of Short-term Gait Adaptation to the Split-belt Treadmill. Neuroscience 451, 36–50.

Hug F, Del Vecchio A, Avrillon S, Farina D & Tucker K (2021). Muscles from the same muscle group do not necessarily share common drive: evidence from the human triceps surae. J Appl Physiol 130, 342–354.

Ivanenko YP, Poppele RE & Lacquaniti F (2004). Five basic muscle activation patterns account for muscle activity during human locomotion. J Physiol 556, 267–282.

Jacobs JV, Wu G & Kelly KM (2015). Evidence for beta corticomuscular coherence during human standing balance: Effects of stance width, vision, and support surface. Neuroscience 298, 1–11.

Jacobsen NA & Ferris DP (2023). Electrocortical activity correlated with locomotor adaptation during split-belt treadmill walking. J Physiol; DOI: 10.1113/JP284505.

Kanazawa H, Kawai M, Kinai T, Iwanaga K, Mima T & Heike T (2014). Cortical muscle control of spontaneous movements in human neonates. Eur J Neurosci 40, 2548–2553.

Kitago T & Krakauer JW (2013). Motor learning principles for neurorehabilitation. Handb Clin Neurol 110, 93–103.

Kristeva R, Patino L & Omlor W (2007). Beta-range cortical motor spectral power and corticomuscular coherence as a mechanism for effective corticospinal interaction during steady-state motor output. Neuroimage 36, 785–792.

Liu J, Sheng Y & Liu H (2019). Corticomuscular Coherence and Its Applications: A Review. Front Hum Neurosci 13, 100.

Maclellan MJ, Ivanenko YP, Massaad F, Bruijn SM, Duysens J & Lacquaniti F (2014). Muscle activation patterns are bilaterally linked during split-belt treadmill walking in humans. J Neurophysiol 111, 1541–1552.

Malone LA, Bastian AJ & Torres-Oviedo G (2012). How does the motor system correct for errors in time and space during locomotor adaptation? J Neurophysiol 108, 672–683.

Mawase F, Haizler T, Bar-Haim S & Karniel A (2013). Kinetic adaptation during locomotion on a split-belt treadmill. J Neurophysiol 109, 2216–2227.

Mendez-Balbuena I, Huethe F, Schulte-Mönting J, Leonhart R, Manjarrez E & Kristeva R (2012). Corticomuscular coherence reflects interindividual differences in the state of the corticomuscular network during low-level static and dynamic forces. Cereb Cortex 22, 628–638.

Mendez-Balbuena I, Naranjo JR, Wang X, Andrykiewicz A, Huethe F, Schulte-Mönting J, Hepp-Reymond M-C & Kristeva R (2013). The strength of the corticospinal coherence depends on the predictability of modulated isometric forces. J Neurophysiol 109, 1579– 1588.

Morton SM & Bastian AJ (2006). Cerebellar contributions to locomotor adaptations during splitbelt treadmill walking. J Neurosci 26, 9107–9116.

Ogawa T, Kawashima N, Ogata T & Nakazawa K (2014). Predictive control of ankle stiffness at heel contact is a key element of locomotor adaptation during split-belt treadmill walking in humans. J Neurophysiol 111, 722–732.

Oshima A, Nakamura Y & Kamibayashi K (2022). Modulation of Muscle Synergies in Lower- Limb Muscles Associated With Split-Belt Locomotor Adaptation. Front Hum Neurosci; DOI: 10.3389/fnhum.2022.852530.

Perez MA, Lundbye-Jensen J & Nielsen JB (2006). Changes in corticospinal drive to spinal motoneurones following visuo-motor skill learning in humans. J Physiol 573, 843–855.

Petersen TH, Kliim-Due M, Farmer SF & Nielsen JB (2010). Childhood development of common drive to a human leg muscle during ankle dorsiflexion and gait. J Physiol 588, 4387–4400.

Petersen TH, Willerslev-Olsen M, Conway BA & Nielsen JB (2012). The motor cortex drives the muscles during walking in human subjects. J Physiol 590, 2443–2452.

Refy O, Blanchard B, Miller-Peterson A, Dalrymple AN, Bedoy EH, Zaripova A, Motaghedi N, Mo O, Panthangi S, Reinhart A, Torres-Oviedo G, Geyer H & Weber DJ (2023). Dynamic spinal reflex adaptation during locomotor adaptation. J Neurophysiol 130, 1008–1014.

Reisman DS, Bastian AJ & Morton SM (2010). Neurophysiologic and rehabilitation insights from the split-belt and other locomotor adaptation paradigms. Phys Ther 90, 187–195.

Reisman DS, Block HJ & Bastian AJ (2005). Interlimb coordination during locomotion: what can be adapted and stored? J Neurophysiol 94, 2403–2415.

Reisman DS, McLean H, Keller J, Danks KA & Bastian AJ (2013). Repeated split-belt treadmill training improves poststroke step length asymmetry. Neurorehabil Neural Repair 27, 460–468.

Riddle CN & Baker SN (2006). Digit displacement, not object compliance, underlies task dependent modulations in human corticomuscular coherence. Neuroimage 33, 618–627.

Roeder L, Boonstra TW & Kerr GK (2020). Corticomuscular control of walking in older people and people with Parkinson’s disease. Sci Rep 10, 2980.

Roeder L, Boonstra TW, Smith SS & Kerr GK (2018). Dynamics of corticospinal motor control during overground and treadmill walking in humans. J Neurophysiol 120, 1017–1031.

Rothwell J, Antal A, Burke D, Carlsen A, Georgiev D, Jahanshahi M, Sternad D, Valls-Solé J & Ziemann U (2021). Central nervous system physiology. Clin Neurophysiol 132, 3043– 3083.

Sato S & Choi JT (2019). Increased intramuscular coherence is associated with temporal gait symmetry during split-belt locomotor adaptation. J Neurophysiol 122, 1097–1109.

Severini G & Zych M (2022). Locomotor adaptations: paradigms, principles and perspectives. Prog Biomed Eng 4, 042003.

Shuman BR, Schwartz MH & Steele KM (2017). Electromyography Data Processing Impacts Muscle Synergies during Gait for Unimpaired Children and Children with Cerebral Palsy. Front Comput Neurosci 11, 50.

Spedden ME, Choi JT, Nielsen JB & Geertsen SS (2019). Corticospinal control of normal and visually guided gait in healthy older and younger adults. Neurobiol Aging 78, 29–41.

Studnicki A & Ferris DP (2023). Parieto-Occipital Electrocortical Dynamics during Real-World Table Tennis. eNeuro; DOI: 10.1523/ENEURO.0463-22.2023.

Studnicki A, Seidler RD & Ferris DP (2023). A table tennis serve versus rally hit elicits differential hemispheric electrocortical power fluctuations. J Neurophysiol 130, 1444– 1456.

Tyrell CM, Helm E & Reisman DS (2014). Learning the spatial features of a locomotor task is slowed after stroke. J Neurophysiol 112, 480–489.

Ushiyama J, Takahashi Y & Ushiba J (2010). Muscle dependency of corticomuscular coherence in upper and lower limb muscles and training-related alterations in ballet dancers and weightlifters. J Appl Physiol 109, 1086–1095.

Ushiyama J, Yamada J, Liu M & Ushiba J (2017). Individual difference in β-band corticomuscular coherence and its relation to force steadiness during isometric voluntary ankle dorsiflexion in healthy humans. Clin Neurophysiol 128, 303–311.

Vargha A & Delaney HD (2000). A Critique and Improvement of the CL Common Language Effect Size Statistics of McGraw and Wong. J Educ Behav Stat 25, 101–132.

Vasudevan EVL & Bastian AJ (2010). Split-belt treadmill adaptation shows different functional networks for fast and slow human walking. J Neurophysiol 103, 183–191.

Yokoyama H, Kaneko N, Masugi Y, Ogawa T, Watanabe K & Nakazawa K (2020). Gait-phase- dependent and gait-phase-independent cortical activity across multiple regions involved in voluntary gait modifications in humans. Eur J Neurosci; DOI: 10.1111/ejn.14867.

Yokoyama H, Kaneko N, Ogawa T, Kawashima N, Watanabe K & Nakazawa K (2019). Cortical Correlates of Locomotor Muscle Synergy Activation in Humans: An Electroencephalographic Decoding Study. iScience 15, 623–639.

Yokoyama H, Yoshida T, Zabjek K, Chen R & Masani K (2020). Defective corticomuscular connectivity during walking in patients with Parkinson’s disease. J Neurophysiol 124, 1399–1414.

